# Mate choice against gene drives leads to evolutionary rescue

**DOI:** 10.1101/2025.02.28.640836

**Authors:** Marilena De Almeida, Jonathan M. Henshaw, Adam G. Jones, Xiaoyan Long

## Abstract

Artificial gene drives offer a promising approach for controlling populations of agricultural pests, invasive species and disease vectors, such as malaria-carrying mosquitoes. Gene-drive alleles induce a transmission bias, meaning there is a greater than 50% probability that they will be inherited by offspring, facilitating their rapid propagation through populations. When these alleles induce high fitness costs, their spread can lead to the eradication of target populations. While gene drives hold great potential, there are various barriers that can hinder their effectiveness. One such barrier, precopulatory mate choice, has been suggested to impact the spread of gene drives, yet it remains underexplored. Here, we use individual-based simulations to investigate the spread of gene drives while allowing female mating preferences to evolve. In our model, gene-drive alleles exhibit transmission bias but also incur survival costs. Crucially, females are assumed to be able to distinguish between drive carriers and wild-type males based on drive-associated phenotypic differences. Consequently, females can evolve to prefer wild-type or drive-carrying males as mates. Our simulations show that under certain parameter settings, mating preferences against drive carriers can evolve during the spread of a gene drive. This can result in the eradication of the drive alleles and the evolutionary rescue of the population. The impact of mate choice was most pronounced when the gene drive spread relatively slowly, as this allowed more time for preferences against drives to evolve. In addition, evolutionary rescue occurred less frequently for recessive drives than for dominant ones. Our results demonstrate that mate choice can indeed impair the effectives of gene drive as a mechanism of population control and should therefore be seen as a potential risk. We further consider the implications for scientists developing gene drives as a population control technology.

**Lay summary:** Most genes exist in multiple versions called alleles, with each having a 50% chance of being passed on by parents during sexual reproduction. However, some alleles, known as gene drives, manipulate inheritance to spread more widely across populations, even if they reduce the fitness of individuals carrying them. Gene drives occur naturally but can also be engineered in the laboratory. Recent technological advancements have led scientists to explore gene drives as a promising tool for controlling pests, invasive species, and disease vectors like malaria-carrying mosquitoes. However, before releasing any gene drives into the wild, it is crucial to assess their feasibility and potential risks. Here, we used computer simulations to explore how artificial gene drives spread when mate choice is allowed to evolve, assuming females can differentiate between males carrying gene drives and those without them. While gene drives have a transmission advantage, they also reduce the survival of carriers. Our results show that evolution can act rapidly after a gene drive is released. Females evolve to discriminate against drive-carrying males, which can sometimes result in the eradication of the gene drive. When gene drives have high survival costs, this evolutionary response can prevent population extinction, potentially undermining the drive’s purpose. The impact of mate choice is greatest when gene drives spread slowly, giving more time for evolution to act. These findings highlight the importance of considering mate choice dynamics when developing gene drives for population control.

## Introduction

Gene drives are genetic elements that bias inheritance in their favour, enabling them to spread rapidly through a population (Burt, 2003; Burt & Trivers, 2006; Wedell et al., 2019). In contrast to Mendelian inheritance, where each allele at a given locus has an equal chance of 50% of being inherited, gene drives can substantially increase the transmission rate of linked alleles (Grunwald et al., 2019; Rode et al., 2019). They achieve this through various mechanisms (Burt, 2014). For example, some gene drives intervene in the meiotic process (e.g., via sperm killing) to ensure they are over-represented in eggs or sperm (Lindholm et al., 2016; Lyttle, 1993; Sandler & Novitski, 1957). Such meiotic drives were originally discovered in natural populations, such as sex ratio distortion in *Drosophila* (Jaenike, 1996) and the *t* complex in house mice *Mus musculus* (Silver, 1993). Other gene drives, such as homing endonuclease genes (HEGs), effectively convert a heterozygote into a homozygote, resulting in both chromosomes acquiring a copy of the HEGs (Burt & Koufopanou, 2004; Chevalier & Stoddard, 2001). HEGs have been discovered in a variety of organisms, including bacteria, fungi, and plants (Stoddard, 2005). Since the development of CRISPR/Cas9 technology (Hsu et al., 2014; Ran et al., 2013), gene drives can now be artificially created and linked to specific “cargo” alleles chosen by researchers (Bassett et al., 2013; Esvelt et al., 2014). When the gene drive is included in the engineered sequences, the modified genes exhibit a transmission bias, allowing them to be favourably passed on to future generations (Burt, 2003).

Artificial gene drives based on the CRISPR/Cas9 system are currently under active development as tools for managing natural populations (Gantz et al., 2015; Hammond et al., 2016; Kyrou et al., 2018). Laboratory studies have shown that CRISPR-based gene drives can spread within a few generations through unstructured populations of several hundreds to thousands of individuals (Gantz et al., 2015; Gantz & Bier, 2015; Hammond et al., 2016; Kyrou et al., 2018; Wang & Jacobs-Lorena, 2013). Theoretical work has further shown that gene-drive propagation can be successful even when there are fitness costs associated with gene drives (Birand et al., 2022; Boëte et al., 2014; Verma et al., 2023). Due to the dual characteristics of high transmission bias but reduced fitness, proposed applications for CRISPR/Cas9 gene drives include (1) reducing or eradicating populations of disease vectors, agricultural pests, or invasive species, for example by propagating alleles that induce sterility (Birand et al., 2022; Kyrou et al., 2018; Simoni et al., 2020; Yadav et al., 2023), and (2) spreading disease-resistance alleles through vector populations (Dong et al., 2018; Gantz et al., 2015; Hoermann et al., 2022). Artificial gene drives have been developed and tested in laboratory populations of several important pests, including the global fruit pests *Drosophila suzukii* (Yadav et al., 2023) and *Ceratitis capitata* (Meccariello et al., 2024), as well as in disease-vectoring mosquitoes such as *Aedes aegypti*, which transmits dengue and Zika (Anderson et al., 2023; Li et al., 2020), and *Anopheles* mosquitoes, the primary vectors of malaria (Box 1; Gantz et al., 2015; Hammond et al., 2016; Kyrou et al., 2018). However, these artificial gene drives are still in the experimental phase. Before releasing such organisms into the wild, it is crucial to study both the feasibility and potential risks of using gene drives as a tool of population control.

#### Box 1: Gene drives for control of malaria

Vector-borne diseases such as malaria, dengue and Zika remain a major public health threat (WHO, 2020). Despite advances in medication and vaccination, malaria alone caused about half a million deaths in 2022 (WHO, 2023). Consequently, innovative intervention strategies are imperative to address the challenges posed by these diseases. To reduce the incidence of malaria using artificially engineered gene drives, two strategies have been developed (Boëte et al., 2014; Burt, 2014; Rode et al., 2019). The first strategy aims to reduce egg production to the point of total population collapse by linking a gene drive to an allele that causes sterility in homozygous females (Kyrou et al., 2018; Rode et al., 2019). Due to their high fitness costs, such sterility alleles would not prevail in natural populations in the absence of a gene drive (Rode et al., 2019). The second strategy is to introduce malaria-refractory alleles into mosquito populations to replace the wild type and reduce the transmission of malaria (Boëte *et al*. 2014; Gantz et al. 2015). Although malaria-refractory alleles are less severe in their fitness costs, laboratory experiments have nonetheless shown that these alleles could not reach fixation without being linked to a drive (Lambrechts et al., 2008). For both strategies – eradication and replacement – transgenic mosquitoes have already been developed, but have not (yet) been released into natural populations (Carballar-Lejarazú et al., 2020; Gantz et al., 2015; Hammond et al., 2016; Kyrou et al., 2018).

Despite their significant potential, gene drives face various barriers that can limit their efficiency. One of the most critical challenges is the evolution of resistance (Champer et al., 2017). Gene drives designed to impose high fitness costs, such as those intended for population eradication, can lead to the selection of alleles that confer resistance to the gene drive (Champer et al., 2017; Drury et al., 2017; Price et al., 2020; Unckless et al., 2017). This resistance can arise quickly, particularly when the fitness costs are severe, thereby constraining or even preventing the dissemination of CRISPR/Cas9-based gene drives (Drury et al., 2017). In addition, population structure and mating systems have been suggested to be crucial determinants of the fixation time and probability of gene drives (Birand et al., 2022; Boëte et al., 2014; Bull et al., 2019; Lindholm et al., 2016; Rode et al., 2019; Verma et al., 2023; Wedell, 2013). In highly sub-structured populations, where most matings happen locally, an eradication drive can lead to local eradication and, therefore, the loss of the drive before it can spread widely (Birand et al., 2022; Bull et al., 2019; Drury et al., 2017). Similarly, in polyandrous species, where females mate with multiple males, competition between the sperm of gene-drive-carrying and wild-type males can prevent offspring from inheriting costly drives, thereby limiting their spread (Manser et al., 2020; Birand et al., 2022). Moreover, the release threshold, which is the minimum number of gene-drive carriers required to successfully invade and spread through a population, plays a crucial role. If this threshold is not met, the gene drive may fail to establish itself, diminishing its effectiveness (Moro et al., 2018; Verma et al., 2023).

Precopulatory mate choice can also impact the spread of gene drives (Boëte et al., 2014; Verma et al., 2023). Since gene drives are often associated with substantial fitness costs, it has been suggested that mating preferences for drive-free partners can evolve when individuals can distinguish between drive carriers and non-carriers at the precopulatory stage (Lande & Wilkinson, 1999; Manser et al., 2017). Such mate choice has also been observed in natural systems (Wedell, 2013). For instance, in stalk-eyed flies, females prefer males with long eye- stalks, which are linked to a genetic suppressor of the drive, over males with short eye-stalks, which are associated with sex-ratio distorters (Cotton et al., 2014; Wilkinson et al., 1998). Another potential example is observed in house mice, where both sexes appear to use olfactory cues to avoid mating with heterozygotes carrying the *t* complex (Lenington, 1991; Price & Wedell, 2008), although this finding has not been definitively confirmed (Sutter & Lindholm, 2016). Precopulatory mate choice might be rare in nature as a strategy to avoid mating with drive carriers, since it requires a strong association between the gene drive and conspicuous traits subject to mate choice (Lande & Wilkinson, 1999; Manser et al., 2017). In contrast, in the context of artificial drives, such association is plausible for two reasons. First, the severe fitness reduction caused by the drive may closely correlate with other phenotypic changes that are perceptible to potential mating partners. Second, drive carriers may be easily identifiable if scientists deliberately link the drive locus to phenotypic markers in order to facilitate tracking the drive’s spread.

Given the potential effect of precopulatory mate choice on the spread of gene drives, and ultimately on population control, it is surprising that very few studies have investigated the theoretical implications of mate choice on population dynamics involving CRISPR-based gene drives. In previous theoretical studies that address this gap, researchers assumed a fixed parameter for mate choice (Boëte et al., 2014; Verma et al., 2023). As a result, it remains unclear whether mate choice can evolve when CRISPR-based gene drives are released into a population, particularly given that transmission bias can cause gene drives to spread rapidly, leading to a lack of variation in traits. To address this, we employ individual-based evolutionary simulations where mate choice and the gene-drive locus coevolve, investigating the conditions under which mate choice can occur and subsequently affect the spread of artificial gene drives. Our goal is to identify potential obstacles to drive-based population control strategies and suggest possible solutions.

### The model

We constructed an individual-based simulation model to study the spread of a gene drive under evolving mating preferences. In our model (Fig. 1), we assumed non-overlapping generations with a single mating season per generation. The gene drive is released at a low initial frequency into a population, after which it coevolves with female mating preferences for gene-drive or wild-type males. In every generation, each female randomly encounters a specified number of males and chooses one of them based on her preferences (details below). A key assumption of our study is that the gene drive influences the phenotype of male carriers, allowing females to distinguish between gene-drive carriers and wild types. After mating, females produce offspring that inherit gene-drive alleles with a fixed transmission bias and preference alleles following Mendelian inheritance principles. The gene drives are assumed to incur fitness costs, reducing the survival probability of offspring. Only surviving offspring become parents, forming the next generation. We built the simulation model using R version 4.2.2 (R Core Team, 2022).

**Figure 1.**
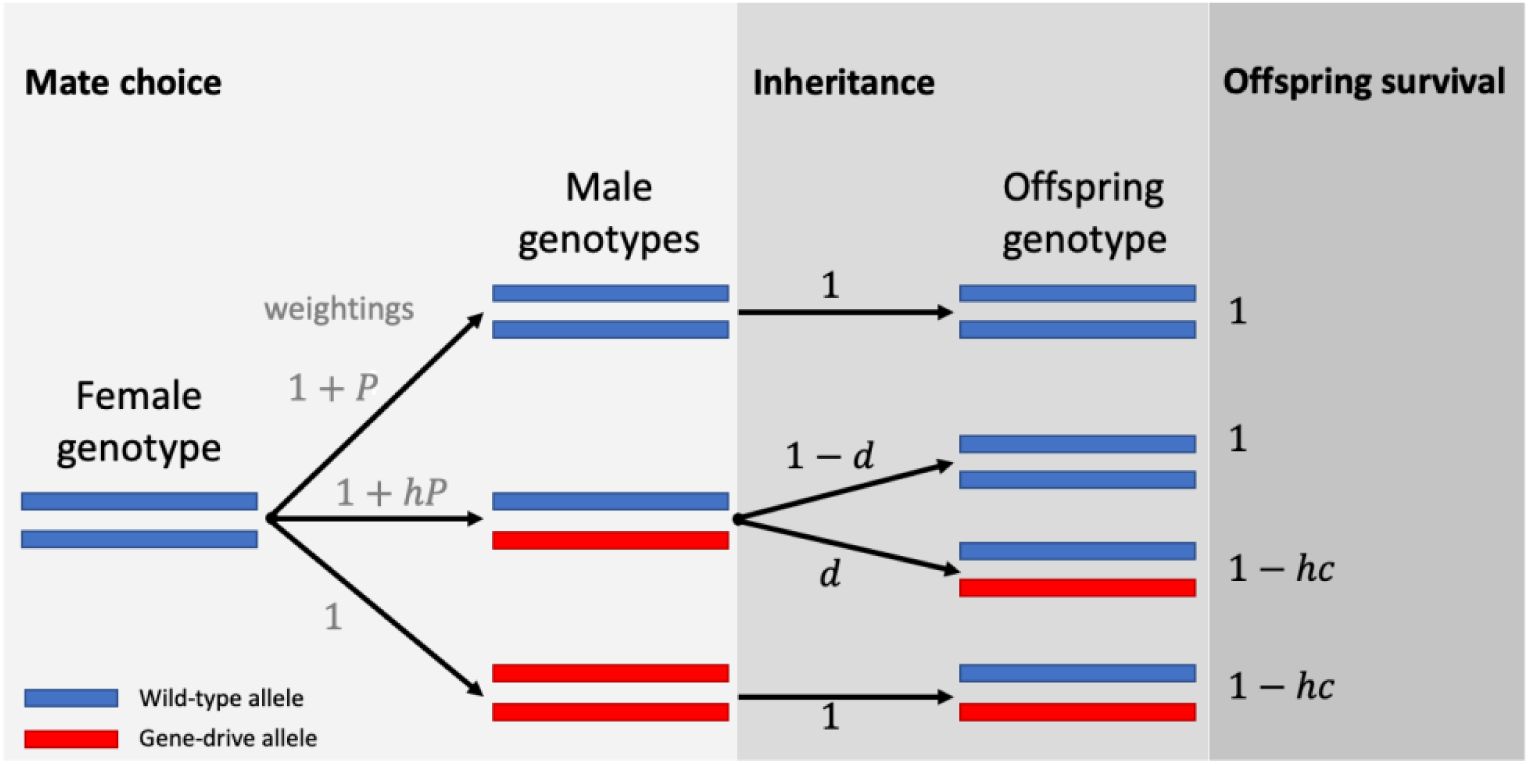
Mate choice, inheritance and offspring survival in our model. Our model considers sexually reproducing populations with a diploid genetic system. There are two genetic loci: The first determines the presence of a gene drive and has two possible alleles: the gene-drive allele (red) and the wild-type allele (blue). The dominance level ℎ determines the phenotype of heterozygotes, with the gene drive being completely dominant when ℎ = 1. The second locus (not shown in the figure) determines female mating preferences, with a continuum of possible alleles represented by real numbers. An individual female’s preference 𝑃 is the average of her two preference alleles, 𝑃 = 1/2(𝑃_1_ + 𝑃_2_). Females with positive preference values (𝑃 > 0) prefer wild-type males, while those with negative values (𝑃 < 0) prefer drive carriers. Each generation, each female encounters a random subset of 𝐴 males. In the depicted example, a wild-type female encounters three males (𝐴 = 3), which happen to be a wild-type male, a heterozygote, and a gene-drive homozygote. The female assigns weightings to each encountered male and chooses her mate probabilistically based on these weightings. In the depicted example, the female is assumed to have a positive preference 𝑃, so she assigns the three males weightings of 1 + 𝑃 (wild-type male), 1 + ℎ𝑃 (heterozygote) and 1 (gene-drive homozygote). The female then chooses her mate with probabilities proportional to these weightings. Subsequently, females produce offspring that inherit gene-drive alleles with a fixed transmission bias (𝑑, where we assume 𝑑 > 0.5 throughout) and preference alleles according to Mendelian inheritance principles. The gene-drive allele imposes survival costs on offspring: The probability of survival is 1 for wild-type offspring, 1 − ℎ𝑐 for heterozygous offspring, and 1 − 𝑐 for homozygous drive carriers (which do not arise in the depicted example because the female is assumed to be wild-type). The surviving offspring replace the current generation and become parents in the next generation.

### Genetics

We model a diploid genetic system with two loci. The first locus determines the presence of gene drive and contains two possible alleles: the gene-drive allele *D* and the wild-type allele *W*. The phenotype of heterozygotes at the drive locus depends on the level of dominance *h* of the drive allele. A dominance level of *h* = 1 means that the gene drive is completely dominant and so heterozygotes are phenotypically indistinguishable from homozygous drive carriers. If *h* = 0, the gene drive is recessive and the phenotype of heterozygotes corresponds to that of wild-types, whereas at *h* = 0.5, heterozygotes are exactly intermediate between the two homozygous phenotypes. In the main text, we primarily present simulation outcomes where the gene drive allele is completely dominant (*h = 1*), with the exception of Fig. 6, where the drive allele is recessive (*h = 0*).

**Table 1:** Definition of parameters and variables used in the model and its default values.

| Symbol | Meaning | Default value |
| --- | --- | --- |
| **Parameters** |  |  |
| $A$ | Number of suitors per female | 4 |
| $c$ | Survival cost of the gene drive | |
| $d$ | Drive efficiency | |
| $F$ | Initial frequency of drive allele | 0.01 |
| $h$ | Dominance of the drive | |
| $K$ | Carrying capacity coefficient | 10 000 |
| $m_D$ | Probability of mutation for the gene-drive allele (per allele per generation) | 0.0001 |
| $m_P$ | Probability of mutation for the mating preference allele (per allele per generation) | 0.0001 |
| $\mu_0$ | Initial mean of the mating preference alleles | 0 |
| $\mu_P$ | Mean mutational effect for the mating preference allele | 0 |
| $N_0$ | Initial population size | 10 000 |
| $\sigma_0$ | Initial standard deviation of the mating preference alleles | 1 |
| $\sigma_P$ | Standard deviation of mutational effects for the mating preference allele | 0.01 |
| $r$ | Maximum growth rate of population | 0.2 |
| $s_{DD}$ | Survival probability of homozygote drive carriers | |
| $s_{WD}$ | Survival probability of heterozygotes | |
| $s_{WW}$ | Survival probability of wild-types | 1 |
| **Variables** |  |  |
| $\lambda$ | Expected number of offspring produced by a female | |
| $N$ | Population size | |
| $P$ | Mating preference | |
| $V_{++}$ | Suitor weighting for homozygotes of the preferred type | |
| $V_{+-}$ or $V_{-+}$ | Suitor weighting for heterozygotes | |
| $V_{--}$ | Suitor weighting for homozygotes of the less preferred type | 1 |

The second locus determines mating preferences, with a continuum of possible alleles represented by real numbers. Mating preferences are only expressed in females. An individual female’s preference value is given by the average of her two preference alleles, 𝑃_1_ and 𝑃_2_:

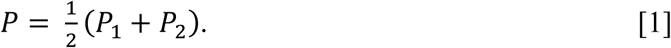

Females with positive preferences (*P* > 0) prefer mating with wild-type males, whereas those with negative preferences (*P* < 0) prefer mating with drive carriers. Females with *P* = 0 show no preference for either phenotype (see next section).

### Mate choice

In every generation, each female mates with a single male. Females choose their mating partners from a pool of *A* suitors. The suitors are drawn uniformly at random with replacement from among the surviving males. Note that the same male could be drawn more than once. Each female prefers either wild-type males or drive carriers, depending on her mating preference *P*. Homozygous suitors of the preferred type (i.e., *WW* if *P* > 0 or *DD* if *P* < 0) are assigned a “weighting” of:

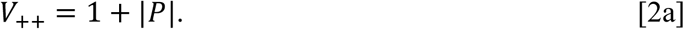

Homozygous suitors of the less-preferred type receive a weighting of:

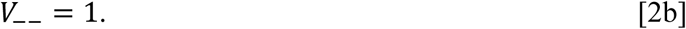

For heterozygote males (i.e., genotype *WD*), weightings are assigned dependent on the dominance level *h* of the drive. Females that prefer drive carriers (*P* < 0) assign heterozygote suitors a weighting of:

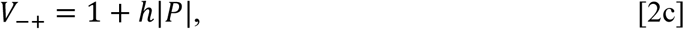

whereas females with a preference for wild-type males (*P* > 0) assign heterozygotes a weighting of:

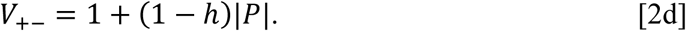

The female chooses her mate from among her suitors with probabilities proportional to these weightings. For example, suppose a female with a mating preference of 𝑃 = 1 is offered three suitors representing the three different genotypes (*WW*, *WD*, *DD*) and the drive allele is partially dominant (ℎ = 0.7). Her weightings are given by 𝑉_++_ = 2 (for the *WW* male), 𝑉_+−_ = 1.7 (for *WD*) and 𝑉_−−_ = 1 (for *DD*). The probability that she mates with the wild-type male is then given by 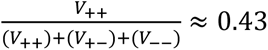 (and approximately 0.36 for the *WD* male and 0.21 for the (𝑉_++_)+(𝑉_+−_)+(𝑉_−−_) *DD* male).

### Population size

Population size is regulated by the interplay between births and survival. We assume that the number of offspring per female is density dependent. For each female, offspring number is drawn from a Poisson distribution with mean given by:

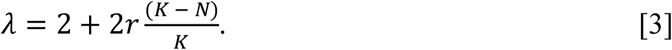

Here *K* is a coefficient that helps determines the population’s carrying capacity, *N* is the current population size and *r* is the maximum rate of population growth. To account for the potential cost for carrying the gene drive, we assume a reduced survival rate for offspring that inherit the gene drive (see next section). In the absence of the gene drive in the population, the population size stabilizes at a size of 𝐾 individuals, at which each female produces an average of two surviving offspring. If drive carriers make up a substantial proportion of the population and the survival cost is high, then the average number of surviving offspring can fall below two per female, leading to a decline in population size. We allowed the population to decline rather than fixing it at the pre-drive carrying capacity, as we are specifically interested in understanding how the coevolution of mate choice and gene-drive spread affects population dynamics.

### Survival costs of the gene drive

The three genotypes (*WW, WD, DD*) have different survival probabilities due to the survival costs of the drive cargo. In our study, we vary these survival costs to understand how they influence the spread and effectiveness of the gene drive within the population. We assume a baseline survival probability of 1 for wild-types and a survival cost of *c* for homozygote drive carriers, yielding survival probabilities of:

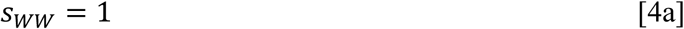

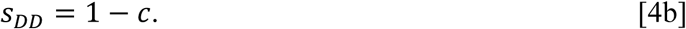

Heterozygote survival depends on the drive dominance level *h* (see above), as follows:

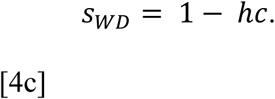

Surviving offspring become the parents of the next generation.

### Inheritance

We allow for a transmission bias in the inheritance of the gene-drive allele. We denote the drive efficiency by *d*, which represents the probability that heterozygotes for the gene-drive allele pass this allele on to any individual offspring. When *d* = 1, the gene-drive is transmitted to all offspring, whereas *d* = 0.5 indicates a 50 % transmission rate, equivalent to unbiased Mendelian inheritance. Throughout our study, we assume 0.5 < 𝑑 ≤ 1, so that the gene-drive allele is always characterized by a transmission bias. For completeness, we also allow for the possibility that drive alleles mutate back to the wild-type allele with a probability of 𝑚_D_ per gene-drive allele per generation. Since this mutation is expected to occur very rarely, if at all, we assume a very low mutation rate. We anticipate that this will not affect our results and conclusions. The mating preference alleles are passed on to the next generation according to standard Mendelian inheritance (Mendel, 1865). Mutational effects at the mating preference locus are normally distributed with mean 𝜇_𝑃_ and standard deviation 𝜎_𝑃_. They occur with a probability of 𝑚_𝑃_ per allele per generation. We assume perfect recombination between the two loci. Offspring were determined to be male or female with equal probability.

### Simulations with and without evolving mating preferences

To better understand how mating preferences evolve and how these preferences affect the spread of the gene drive within population, we compared simulations under two different scenarios: one with evolving mating preferences and one without. In the absence of evolving mating preferences, mating was assumed to be random with respect to the drive locus. In this case, the preference genes play no role in partner choice. In both cases, we considered a broad range of survival costs associated with carrying the gene drive and the transmission bias of the drive allele.

### Initialization

We initialized simulation runs with a population of 10 000 individuals. The gene drive was released once into the population at the beginning of a simulation run, with 𝐹 being the initial probability for an allele to be a drive allele. Mating preference alleles were sampled out of a normal distribution with mean 𝜇_0_and standard deviation 𝜎_0_. Half of the initial population was assigned to be female, the other half as male. Parameters and their default values are summarized in Table 1. For most parameter combinations presented in the main text, we conducted 130 replicated simulations (with exceptions for Fig. 6), each running over 200 generations.

## Results

### The gene drive invades when drive efficiency is high relative to survival costs

As previously shown by Rode et al. (2019) and Verma et al. (2023), the invasion potential of a gene drive is dependent on both the gene drive’s transmission efficiency and gene drive- associated costs. Here, we first present three basic scenarios for the spread of a completely dominant gene drive, which each arise under particular combinations of drive efficiency and drive-associated survial costs (Fig. 2). First, when both drive efficiency and the survival probability of drive carriers were high (𝑠_𝐷𝐷_ = 𝑠_𝑊𝐷_ = 0.9, 𝑑 = 0.9: Fig. 2A, note that 𝑠_𝑊𝑊_ = 1 for all scenarios), the gene drive rapidly took over the population (in less than twenty generations on average for our parameter settings, Fig. 2A_1_). Due to the drive’s survival costs, the maximum population size at equilbrium fell as the drive took over the population (Fig. 2A_2_). In the short space of time it took for the gene drive to reach fixation, females did not evolve strong preferences for either type of suitor on average (Fig. 2A_3_).

**Figure 2.**
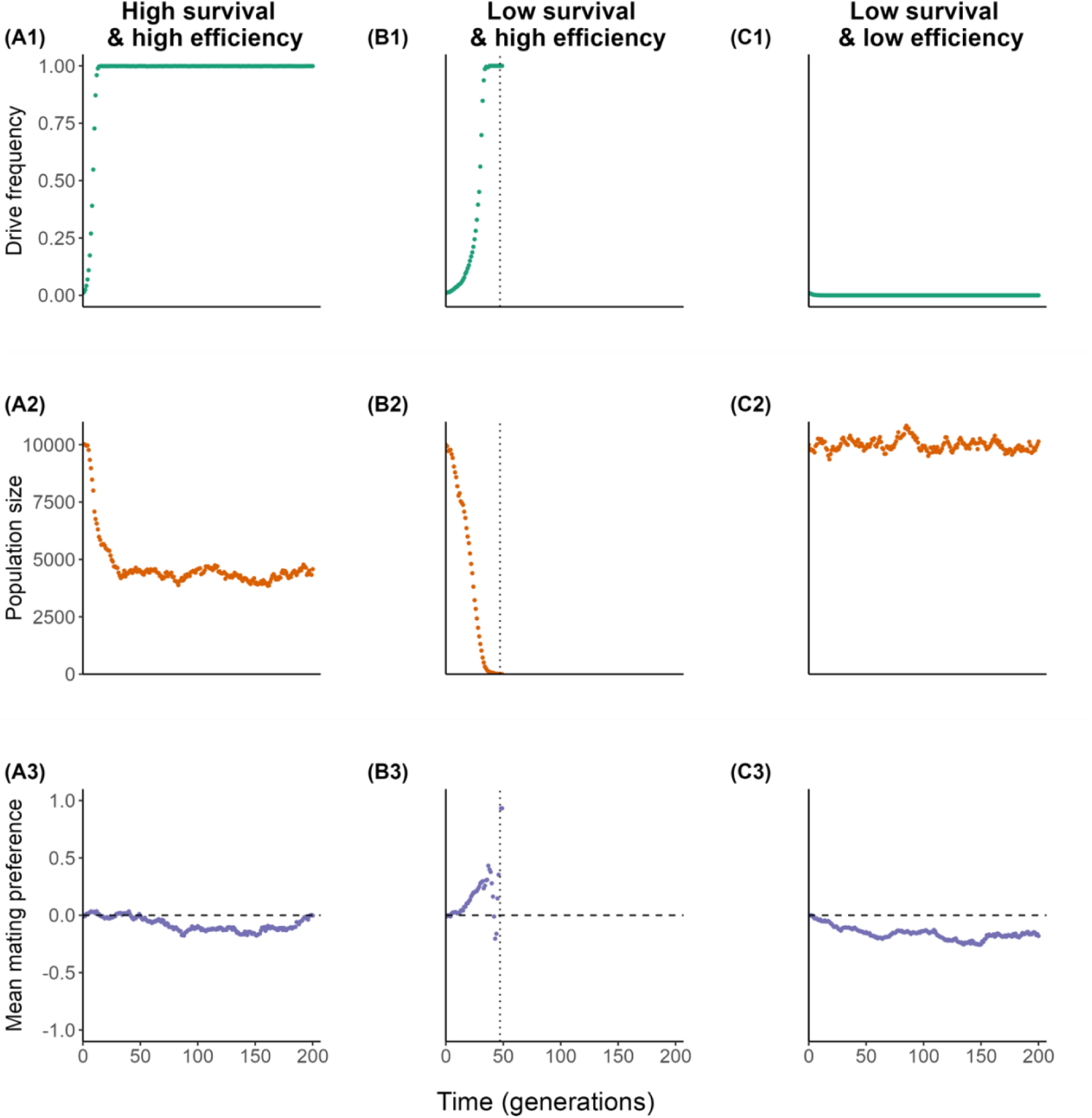
Impacts of drive efficiency and survival costs on drive-allele frequency, population size, and mating preference. This figure illustrates the evolution of **(1)** the drive-allele frequency (green), **(2)** the population size (orange), and **(3)** the mean mating preference (purple) under different levels of drive efficiency and survival costs of gene-drive carriers. Each column (A, B, C) represents a representative simulation replicate selected from 130 replicates for each parameter setting. **A:** High survival of gene-drive carriers and high drive efficiency (𝑠_𝐷𝐷_ = 0.9, *d* = 0.9). **B:** Low survival of gene-drive carriers and high drive efficiency (𝑠_𝐷𝐷_ = 0.6, *d* = 0.9). **C:** Low survival of gene-drive carriers and low drive efficiency (𝑠_𝐷𝐷_ = 0.6, *d* = 0.6). In all scenarios, 𝑠_𝑊𝐷_ = 𝑠_𝐷𝐷_ and 𝑠_𝑊𝑊_ = 1. The vertical black dotted lines in panel B indicate the average time for a population to go extinct across 130 replicates. The horizontal dashed lines in panels A3, B3, and C3 mark a mean female mating preference of zero. All other parameter values are as listed in Table 1.

Second, when high drive efficiency was combined with relatively low survival of drive carriers (𝑠_𝐷𝐷_ = 𝑠_𝑊𝐷_ = 0.6, 𝑑 = 0.9; Fig. 2B), the gene drive still rapidly dominated the population (Fig. 2B_1_), but the high survival costs ultimately resulted in the extinction of the population (Fig. 2B_2_). Before the population went extinct, females evolved to prefer wild-type males (Fig. 2B_3_), but this preference was insufficient to prevent the drive from taking over (cf. Fig. 3). When the population size became very small (in the last few generations before extinction), there was strong drift in mating preferences.

**Figure 3.**
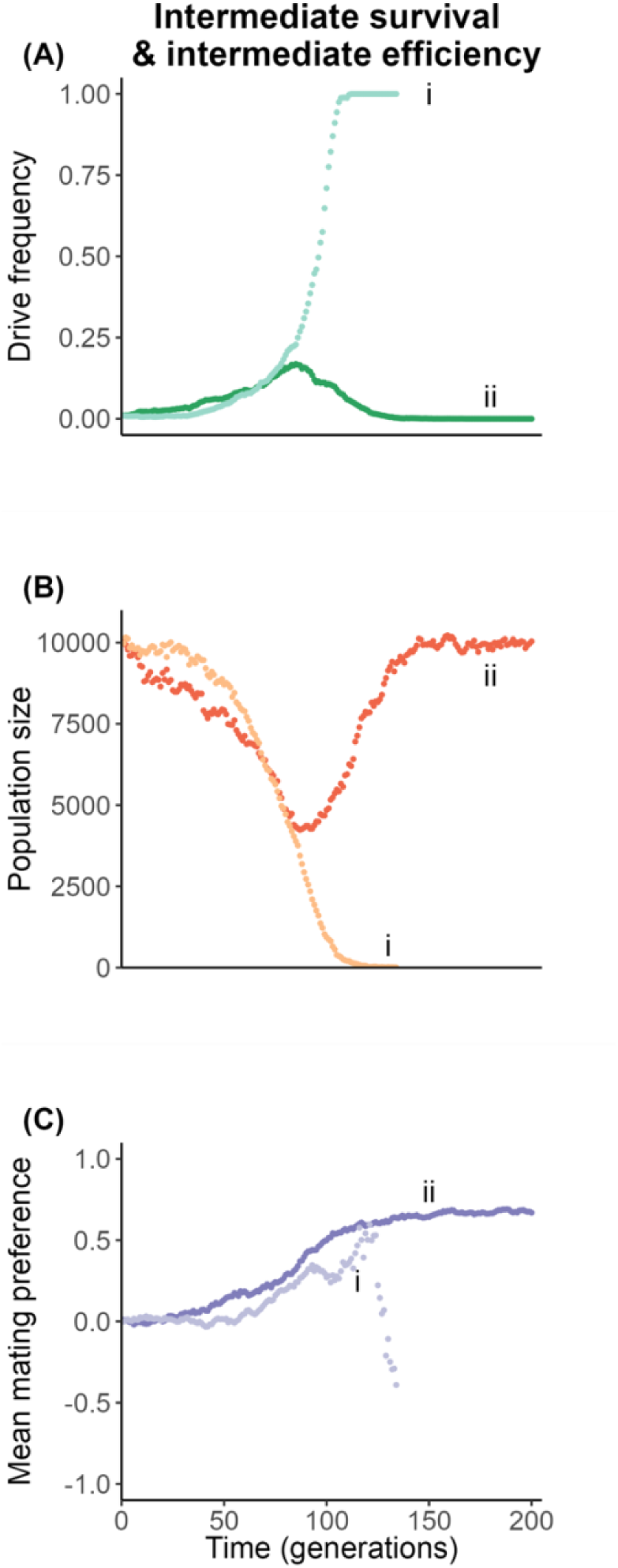
Evolutionary rescue can occur when the population exhibits intermediate levels of drive survival costs and drive efficiency. The figure illustrates the evolution of **(A)** the gene- drive allele frequency, **(B)** population size, and **(C)** mean mating preference. We present two representative replicates (i and ii) from a total of 130 replicates with the same parameter settings. In roughly 30% of the 130 replicate simulations (consistent with replicate i), the gene drive took over the population (light green in A), leading to population extinction (light orange in B), and females developed mating preferences for wild-type males (light purple in C). In 65% of the replicate simulations (as in replicate ii), the gene drive was eliminated from the population (dark green in A) due to strong mating preferences for wild-type males (dark purple in C), resulting in evolutionary rescue of the population (dark orange in B). In one replicate, neither the drive nor the wild-type reached fixation within 200 generations. Here, the survival of drive carriers was set to 𝑠_𝐷𝐷_ = 𝑠_𝑊𝐷_ = 0.7 and drive efficiency to *d* = 0.72. All other parameter values are as listed in Table 1.

Third, simulations with low drive-carrier survival and low drive efficiency resulted in the gene drive being quickly eliminated from the population (𝑠_𝐷𝐷_ = 𝑠_𝑊𝐷_ = 0.6, 𝑑 = 0.6; Fig. 2C_1_). In this case, the population size remained at its original carrying capacity because the number of drive carriers stayed consistently low (Fig. 2C_2_). Females did not evolve strong mating preferences as the gene drive rapidly disappeared from the population, leaving no variation in male traits to choose from. After the gene drive was lost, any evolution of mating preferences was a result of genetic drift (Fig. 2C_3_).

### Sexual selection can impede the spread of the gene drive

When either drive efficiency or the survival of drive carriers was high (Fig. 2), sexual selection sometimes occurred (Fig. 2B), but was insufficiently strong to influence the ultimate fate of the drive allele. In contrast, at intermediate levels of drive survival costs and drive efficiency, the evolution of female preferences for wild-type males could ‘rescue’ the population from drive take-over (Fig. 3). With intermediate drive efficiency and survival (e.g., 𝑠_𝐷𝐷_ = 𝑠_𝑊𝐷_ = 0.7, 𝑑 = 0.72), the simulations resulted in two alternative outcomes (Fig. 3). In one scenario (represented by the lighter-coloured replicate i in Fig. 3), the gene drive rapidly dominated the population (light green in Fig. 3A), resulting in population extinction (light orange in Fig. 3B), accompanied by a short period of female preferences for the wild-type (light purple in Fig. 3C). Given the lower drive survival costs compared to Fig. 2B, the average time until population extinction was prolonged.

In the other scenario (darker-coloured replicate ii in Fig. 3), after an initial increase in drive allele frequency, the gene drive declined until it disappeared entirely (dark green in Fig. 3A). Concurrently with the initial spread of the drive, the population size decreased, but it recovered as the drive diminished, indicating an evolutionary rescue (dark orange in Fig. 3B). This rescue occurred due to the evolution of strong female preferences for wild-type males who have the highest survial probability (dark purple in Fig. 3C).

The occurrence of the two alternative outcomes can be explained as follows. Under these parameter settings, drive efficiency and survival were not high enough for the drive to rapidly reach fixation. There was consequently a relatively long window of opportunity for the evolution of female preferences for the wild type. If female preferences for wild-type males developed to be sufficiently strong before the drive took over the population, then the initial fitness advantage of the drive allele was reversed, leading to an evolutionary rescue. Otherwise, the drive took over and preferences were rendered selectively inert.

### Drive survival costs and drive efficiency shape the potential for evolutionary rescue

Fig. 3 demonstrated that sexual selection can hinder the spread of the gene drive, leading to evolutionary rescue under specific combinations of drive survival costs and efficiency. In Figs. 4 and 5, we systematically investigate the conditions under which sexual selection can eliminate the gene drive and the impact of sexual selection on the probability of population extinction. Consistent with the observations presented in Figs. 2A and 2B, the gene drive rapidly went to fixation when the fitness of the drive allele was high, which was determined by a (linear) combination of drive efficiency and survival of the gene drive (red circles in Fig. 4). When drive carriers had small survival costs, drive take-over resulted in population shrinkage but not extinction (as in Fig. 2A). This is because the additional mortality due to the drive was offset by an increased growth rate as population size decreased. With a maximum population growth rate of *r* = 0.2 in our model, a survival rate as low as 𝑠_𝐷𝐷_ = 𝑠_𝑊𝐷_ = 0.8 for drive carriers could still be borne without resulting in extinction (light yellow squares on the right-hand side of Fig. 5B). When drive survival was low (as in Fig. 2B), its take-over drove all populations to extinction (black squares in Fig. 5B). Conversely, under conditions of low drive efficiency and/or a low survival rate of the gene drive, the gene drive failed to spread through the population and was eventually eradicated (blue circles in Fig. 4). The corresponding evolutionary trajectories were similar to those depicted in Fig. 2C. Due to the elimination of the drive, the extinction probability of the population went to zero (light yellow squares in the bottom-left corner of Fig. 5B).

**Figure 4.**
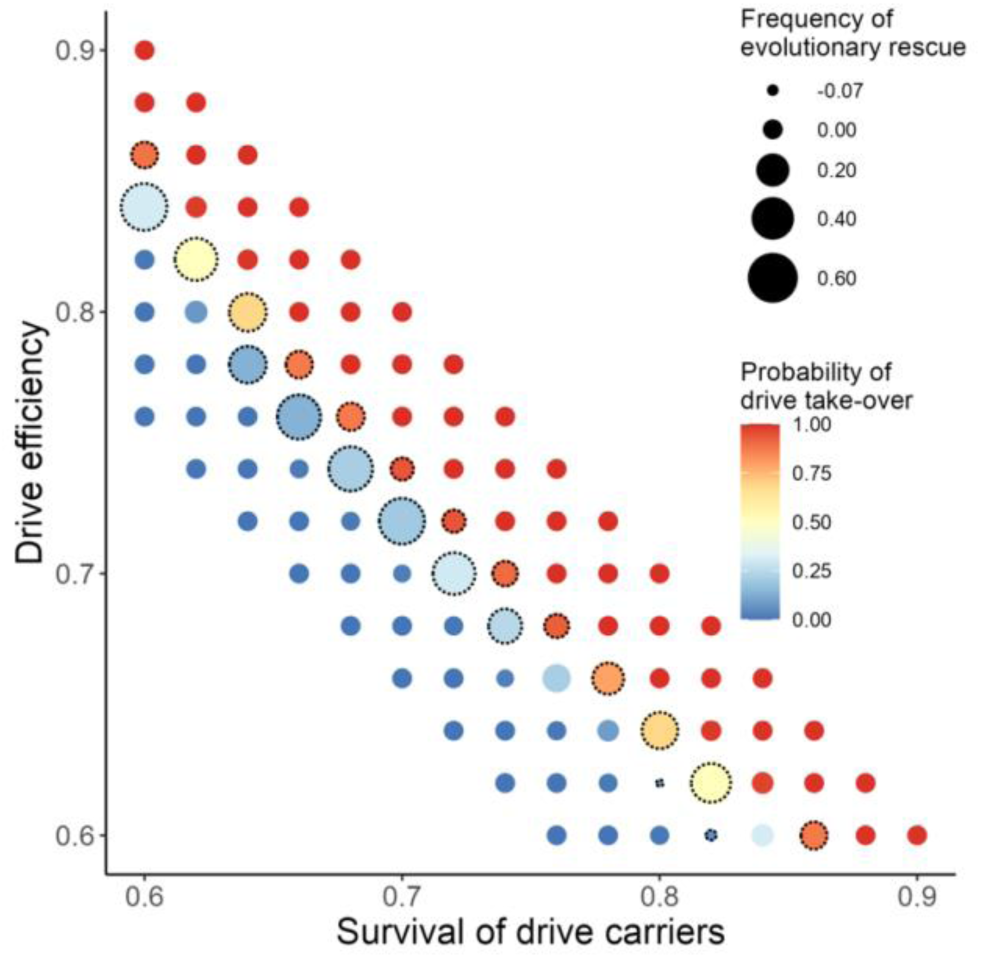
Effects of drive efficiency and survival of drive carriers on the probability of gene drive take-over and the occurrence of evolutionary rescue. The probability of drive take-over is measured as the average frequency of the drive allele at generation 200 in scenarios with evolving mating preferences across 130 replicates (indicated by circle colour). As with Figs. 2 and 3, the gene drive typically either took over the entire population or was completely eliminated within 200 generations. The frequency of evolutionary rescue is calculated as the difference in the probability of drive take-over between simulations that include the evolution of mating preferences and those that do not (i.e., random mating) across 130 replicates per scenario (indicated by circle size; larger circles indicate a higher occurrence of evolutionary rescue due to the evolution of female preferences for wild-type males). The dashed circles indicate a significant difference in the frequency of evolutionary rescue between the two scenarios (Fisher’s exact tests on the proportions of runs with drive allele frequencies > 0.5 in generation 200, 𝑝 < 0.05). The highest frequency of evolutionary rescue via sexual selection (0.58) occurred at 𝑠_𝐷𝐷_ = 𝑠_𝑊𝐷_ = 0.6 and *d* = 0.84, while the lowest frequency (−0.07) was observed at 𝑠_𝐷𝐷_ = 𝑠_𝑊𝐷_ = 0.8 and *d* = 0.62, indicating a higher frequency of the drive in simulations with evolving mate choice compared to those with random mating. No long-term coexistence of the two alleles was observed in any of our simulations. All parameter values are detailed in Table 1. Intriguingly, at high drive-carrier survival rates and low drive efficiencies, there was a limited parameter range where the probability of drive take-over was significantly higher in simulations with mating preferences than in those with random mating (e.g., 𝑠_𝐷𝐷_ = 𝑠_𝑊𝐷_ = 0.78 and 𝑑 = 0.66: Figs. 4,5). This appears to be the result of a curious quirk of our model: If roughly half of females initially prefer wild-type males and the other half prefer gene-drive carriers, then there is a small mating advantage to the rarer phenotype. This small advantage can be sufficient to boost the initial spread of the gene drive, even if females subsequently evolve to prefer wild- type males.

**Figure 5.**
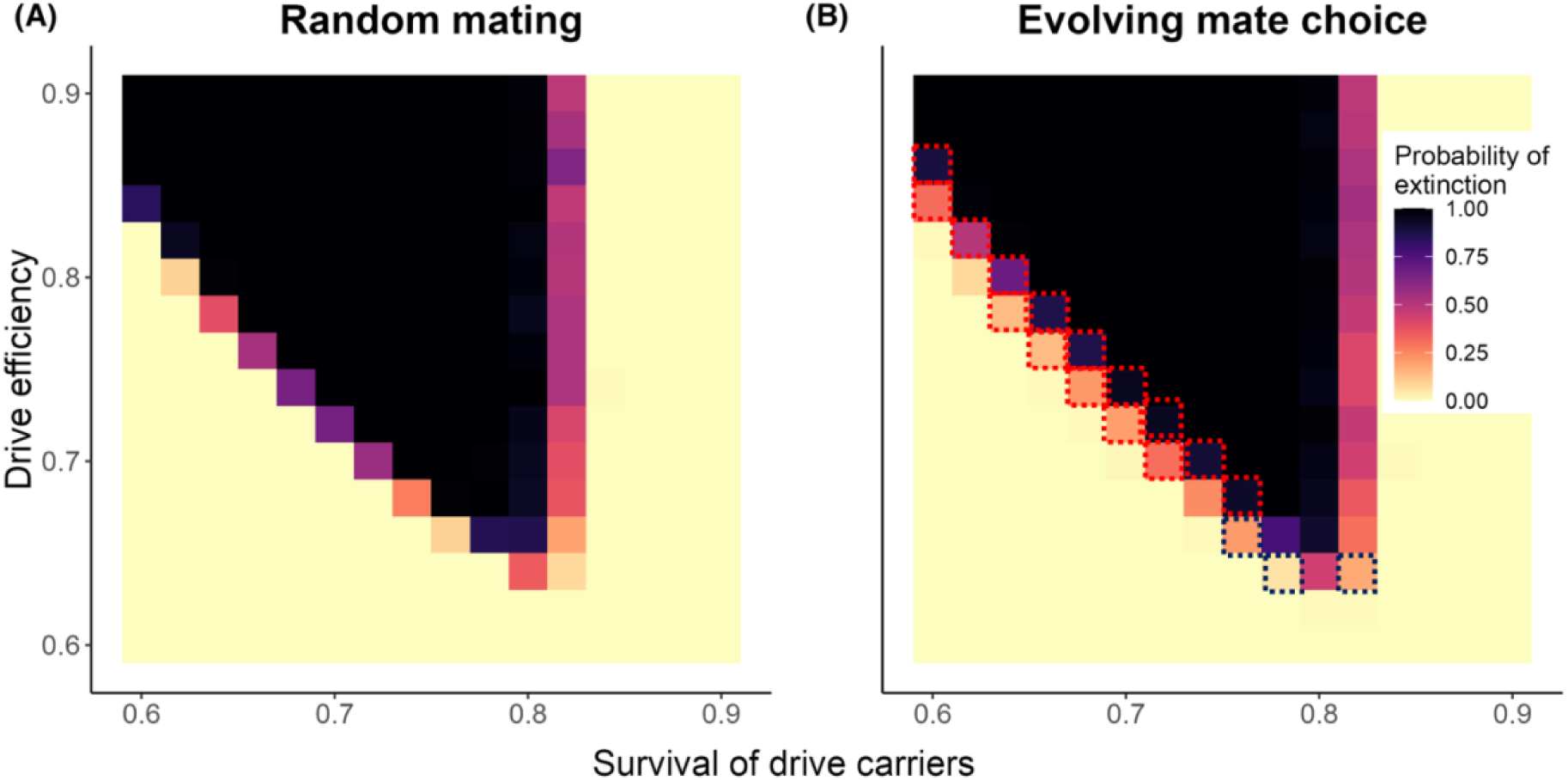
Effects of drive efficiency and survival of drive carriers on the probability of population extinction. The probability of extinction (indicated by square colour) is defined as the frequency of populations that went extinct over the course of 200 generations across 130 replicates per scenario. We compared **(A)** simulations without evolving mating preferences (i.e., random mating) to **(B)** simulations allowing for evolving mate choice. Scenarios where the probability of extinction was significantly lower with evolving mate choice than with random mating are outlined in red, while opposite cases (significantly higher extinction with evolving mate choice) are outlined in dark blue (Fisher’s exact tests on the proportions of runs that resulted in extinction, 𝑝 < 0.05). All other parameters are as detailed in Table 1.

**Figure 6.**
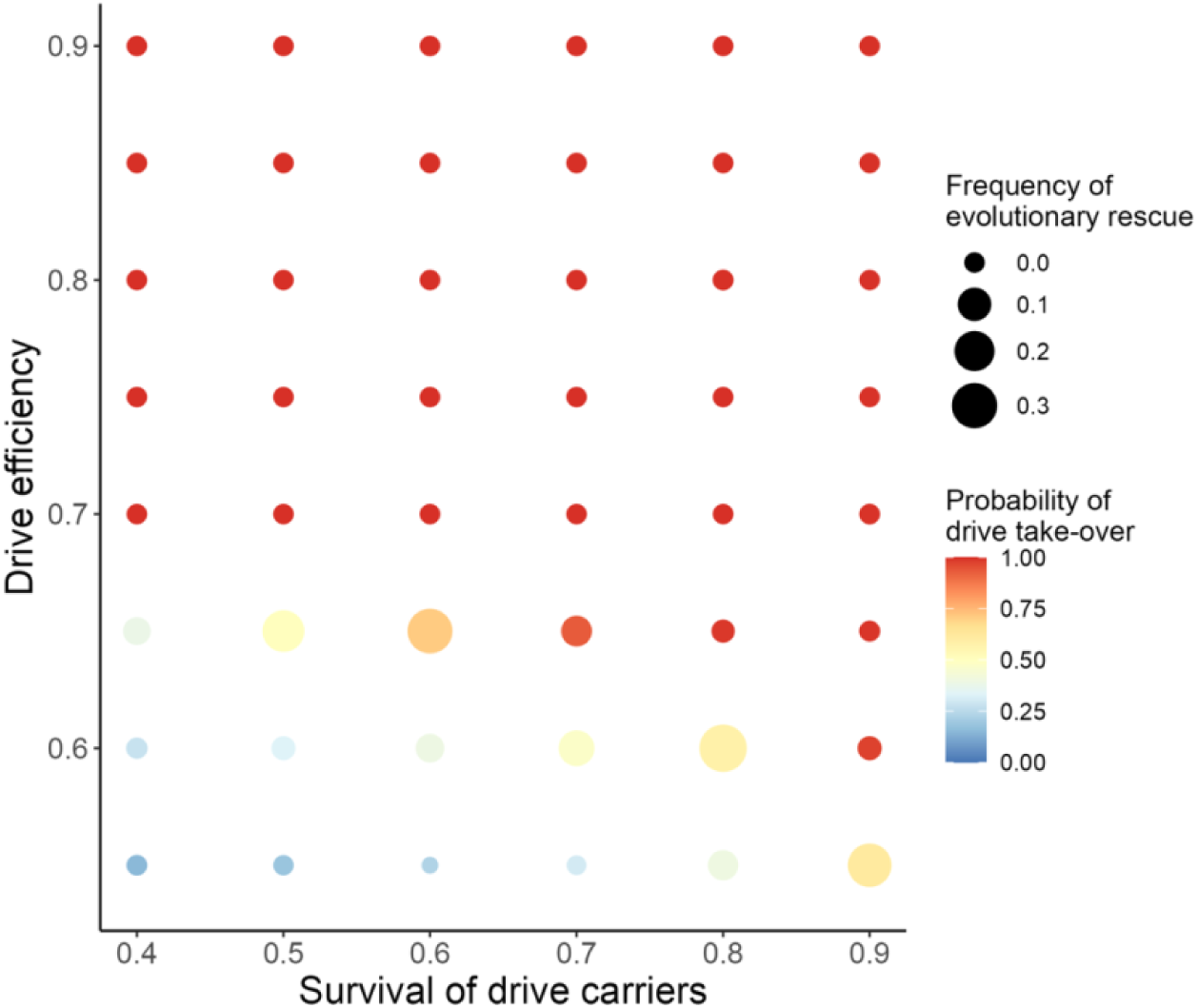
Probability of gene drive take-over and the frequency of evolutionary rescue when the gene drive is fully recessive (*h* = 0). The probability of drive take-over (indicated by circle colour) is measured as the average frequency of the drive allele at generation 200 across 20 replicates. The frequency of evolutionary rescue (indicated by circle size) is calculated as in Fig. 4. All other parameter values are as detailed in Table 1 in the main text.

Within a specific range of drive survival costs and drive efficiency (along the diagonal in Figs. 4 and 5), the drive neither rapidly dominated nor was immediately eliminated from the population. This allowed sufficient time for the evolution of mating preferences for the wild type, often resulting in evolutionary rescue (as in Fig. 3). However, evolutionary rescue could also occur through genetic drift, as the population rapidly declined alongside the spread of the gene drive, making the stochastic loss of the drive possible. To quantify the frequency of evolutionary rescue via sexual selection, we compared the number of simulation replicates resulting in drive take-over between scenarios with evolving mating preferences and those with only random mating (i.e., no evolving mating preferences). For some parameter settings, the drive was eliminated up to 58% more frequently when sexual selection against drive carriers was possible (Fig. 4). As expected, populations with evolving mate choice generally exhibited a higher frequency of evolutionary rescue (Fig. 4) and, thus, lower extinction probabilities (Fig. 5) compared to those with random mating.

### Effects of dominance of the gene-drive allele

Up until now, our analysis has only considered complete dominance of the gene-drive allele (ℎ = 1). We also considered the case where the drive allele was fully recessive (ℎ = 0), so that heterozygotes bore no survival costs (Figs. 6 and S1). In line with the findings of Unckless et al. (2015 & 2017), recessive drives could take over for a wider range of survival and efficiency values than dominant drives (see Fig. 6): With a high drive efficiency, the gene drive always took over the population, independently of the survival costs of homozygote drive carriers (𝑠_𝐷𝐷_ > 0.4, 𝑠_𝑊𝐷_ = 𝑠_𝑊𝑊_ = 1 and *d* = 0.9). Even when the drive efficiency was intermediate (𝑠_𝐷𝐷_ > 0.4, 𝑠_𝑊𝐷_ = 𝑠_𝑊𝑊_ = 1, *d* = 0.7), the gene drive always spread through the population. The gene drive was only lost when low drive efficiency was combined with low survival of homozygote drive carriers (e.g., 𝑠_𝐷𝐷_ = 0.4, 𝑠_𝑊𝐷_ = 𝑠_𝑊𝑊_ = 1, *d* < 0.7). Moreover, evolutionary rescue occurred less frequently for recessive drives (cf. Figs. 4 and 6). Consistent with Rode et al. (2019), our results also show that low drive efficiencies (*d* < 0.7) can lead to the coexistence of the wild-type allele and the recessive gene-drive allele (Fig. S1). For parameter settings where the gene drive would normally take over in the absence of mate choice, the evolution of preferences for wild-types could lead to a decrease in the drive allele frequency, fostering coexistence by preventing the drive from taking over completely (e.g., 𝑠_𝐷𝐷_ = 0.5, 𝑠_𝑊𝐷_ = 𝑠_𝑊𝑊_ = 1, *d* = 0.65).

### Effects of initial conditions of mating preferences

Initial mating preferences had a strong effect on the course and outcomes of the gene drive. First, with much smaller initial variation (𝜎_0_ = 0.1) around the mean preference value of zero, we did not observe any evolutionary rescue scenarios (Fig. S2). Second, if the mean preferences were initially biased towards wild-type males, the gene drive was eliminated from most of the populations (Fig. S3). In this case, drive take-overs only occurred under very high drive efficiency and survival. Conversely, an initial preference for drive carriers led to drive take- over for most survival and efficiency conditions (Fig. S4).

## Discussion

Our simulation model predicts that drive transmission efficiency and drive-associated costs are crucial factors influencing the spread of a gene drive and the development of mate choice. Consistent with previous studies (Rode et al., 2019; Verma et al., 2023), our simulations predict rapid fixation of the drive allele when transmission efficiency is high and associated costs are relatively low (Fig. 4). Drive take-over ultimately led either to population extinction (at high costs, see Figs. 2B and 4) or to a surviving population dominated by the drive allele (see Figs. 2A and 4). In contrast, low drive efficiencies and high costs slowed down or even prevented the drive’s spread (see Figs. 2C and 4). Interestingly, our study shows the potentially constraining role of mate choice in the spread of a gene drive (Figs. 3 and 4). Under conditions where the gene drive could not spread rapidly (e.g., with a low drive efficiency and intermediate drive-associated costs), strong preferences for the wild type evolved, often leading to the extinction of the drive and the evolutionary rescue of the population (Figs. 3 and 4). Overall, we observed less frequent drive take-overs, and consequently fewer population extinctions, in simulations with evolving mating preferences than in those without (Figs. 4 and 5). The evolution of mate choice is most likely when (1) there is sufficient time for mate choice to evolve before the drive goes to fixation, and (2) other alleles (e.g., those underlying mating preferences) are under strong selection to avoid being linked to the drive allele. Both conditions are more easily met when the gene drive imposes a high cost on its bearers.

Our results suggest that the emergence of mate choice should be taken into account when assessing gene drives for release into natural populations. We make three recommendations: (1) For eradication drives, researchers should favor cost mechanisms that are less likely to be detected by potential mating partners. For example, sterility alleles may be a more effective choice than alleles reducing survival, as the latter may have correlated effects on the phenotype of surviving drive-carrying individuals. (2) Researchers should avoid including additional phenotypic markers (e.g., for tracking purposes) in the gene drive’s payload unless absolutely necessary. If included, such markers should be chosen to be imperceptible to the sensory system of the target species. (3) Existing mating preferences for or against drive-carrying individuals should be assessed under laboratory conditions before any release. Our model predicts that existing preferences against gene-drive carriers (Fig. S3), as well as substantial standing variation for such preferences (cf. Fig. S2), make evolutionary rescue more likely. To increase the chances of a successful spread, gene drives could potentially even be coupled with known attractive traits or with preferences for drive-linked traits.

Further, our model showed that recessive drives could take over for a wider range of drive survival and efficiency values, and that evolutionary rescue occurred less frequently for recessive than for dominant drives (Figs. 4 and 6). This is because fitness costs associated with a recessive drive only arise in homozygous drive carriers (Deredec et al., 2008; Unckless et al., 2015 & 2017). Heterozygotes, in contrast, carry the drive silently without bearing any costs. This argument also extends to mate choice if females are unable to distinguish between wild-type individuals and heterozygous drive carriers. Consequently, even when mate choice acts against the spread of the gene drive, recessive drives can still persist at intermediate frequencies in the population (Fig. 6). Therefore, our results support previous studies (Deredec et al., 2008; L. Lehmann et al., 2007; Unckless et al., 2017) that recessive drive offers greater robustness and effectiveness in population control over the long term, although dominant drives might seem like a wise choice to rapidly reduce population size. However, the coexistence of recessive drives and wild types could pose a problem if a drive is introduced into populations through multiple releases. This is because the continued coexistence of the drive allele and the wild-type allele supports the evolution of mate choice (L. Lehmann et al., 2007).

Our model assumes a single panmictic population, wherein population density is regulated logistically to prevent the population size from exceeding 10 000 individuals. However, in certain scenarios, the population experiences a great decline as gene drives carrying survival costs become predominant. This substantial reduction in population size can lead to genetic drift, influencing the evolution of mate choice in our study. To isolate the effects of sexual selection on gene drive spreading and validate the robustness of our simulation predictions, future simulations that minimize variation in the strength of drift—such as maintaining the population at a large size—could be highly beneficial. Moreover, natural populations are often highly structured and may experience strong temporal fluctuations in size. For instance, mosquito populations commonly exceed 10 000 individuals, but can also go through seasonal bottlenecks with only a small fraction of the population surviving (T. Lehmann et al., 1998). Eradication drives may be more effective when released during such bottlenecks, as populations lacking in genetic diversity should be both less able to evolve mate choice and less able to recover from a population crash (Nabutanyi & Wittmann, 2021). Further, theoretical studies have shown that population structure can slow the progress of gene drives (Birand et al., 2022; Verma et al., 2023). This should allow more time for mating preferences to evolve. Hence, in highly structured populations, we should expect evolutionary rescue via mate choice to occur more frequently.

In our model, mating dynamics is highly simplified relative to natural populations. We assume that females choose from among a fixed number of suitors. A higher number of suitors per female should increase intrasexual competition and allow for more efficient mate choice, leading to more evolutionary rescue scenarios. Moreover, our study does not account for the potential costs associated with the evolution of female preferences, which can significantly influence the coevolution of preferences and preferred traits (Manser et al., 2017; Pomiankowski, 1987). Therefore, further research is needed to explore more realistic scenarios, such as the coevolution of female preferences (including associated costs), male traits, and the drive locus, to better understand the interaction between sexual selection and the spread of CRISPR-based gene drives. Last but not least, despite encountering multiple males, females only mate with one of them, thus eliminating the potential for sperm competition among different types of males in our model. Interestingly, in the presence of naturally occurring drives, increased levels of polyandry serve as an important strategy to reduce the likelihood of offspring inheriting costly drives (e.g., in stalk-eyed flies and in house mice, see Lande & Wilkinson, 1999; Manser et al., 2020; Sutter & Lindholm, 2016). Understanding a species’ natural mating system is thus essential for assessing the effectiveness of gene drives as a tool of population control. For future research, it would be valuable to consider both pre- and postcopulatory mate choice to gain a comprehensive understanding of how different mate choice processes, and their interactions, influence gene-drive dynamics.

## Conclusions

We have shown that mate choice can indeed impair the effectives of gene drives as a population control mechanism and should therefore be seen as a potential risk. As we built a highly simplified simulation model to generate our results, it was not possible to capture the full empirical complexity of any gene-drive implementation. Many environmental and ecological factors can influence the spread of a drive (Verma et al., 2023). For malaria-transmitting mosquitoes in particular, further information about the mating system and population structure would be needed for a more realistic risk assessment. However, our model shows that the risk of mate choice evolving against gene drives could be minimized by avoiding the use of phenotypic markers and by releasing recessive rather than dominant drives.

## Author contributions

A.G.J. came up with the original idea. M.A., J.M.H. and X.L. designed the model. M.A. wrote the model code and analysed the results. M.A., J.M.H. and X.L. interpreted the results. M.A. wrote the manuscript. M.A., J.M.H., A.G.J. and X.L. revised the manuscript.

## Data and code availability

R code for the individual-based simulations is available in the Supplementary Materials.

## References

Anderson, M. A. E., Gonzalez, E., Ang, J. X. D., Shackleford, L., Nevard, K., Verkuijl, S. A. N., Edgington, M. P., Harvey-Samuel, T., & Alphey, L. (2023). Closing the gap to effective gene drive in Aedes aegypti by exploiting germline regulatory elements. Nature Communications, 14(1), 338. 10.1038/s41467-023-36029-7

Bassett, A. R., Tibbit, C., Ponting, C. P., & Liu, J.-L. (2013). Highly Efficient Targeted Mutagenesis of Drosophila with the CRISPR/Cas9 System. Cell Reports, 4(1), 220–228. 10.1016/j.celrep.2013.06.020

Birand, A., Cassey, P., Ross, J. V., Russell, J. C., Thomas, P., & Prowse, T. A. A. (2022). Gene drives for vertebrate pest control: Realistic spatial modelling of eradication probabilities and times for island mouse populations. Molecular Ecology, 31(6), 1907–1923. 10.1111/mec.16361

Boëte, C., Agusto, F. B., & Reeves, R. G. (2014). Impact of mating behaviour on the success of malaria control through a single inundative release of transgenic mosquitoes. Journal of Theoretical Biology, 347, 33–43. 10.1016/j.jtbi.2014.01.010

Bull, J. J., Remien, C. H., & Krone, S. M. (2019). Gene-drive-mediated extinction is thwarted by population structure and evolution of sib mating. *Evolution*, Medicine, and Public Health, 2019(1), 66–81. 10.1093/emph/eoz014

Burt, A. (2003). Site-specific selfish genes as tools for the control and genetic engineering of natural populations. Proceedings of the Royal Society of London. Series B: Biological Sciences, 270(1518), 921–928. 10.1098/rspb.2002.2319

Burt, A. (2014). Heritable strategies for controlling insect vectors of disease. Philosophical Transactions of the Royal Society B: Biological Sciences, 369(1645), 20130432. 10.1098/rstb.2013.0432

Burt, A., & Koufopanou, V. (2004). Homing endonuclease genes: The rise and fall and rise again of a selfish element. 14. https://linkinghub.elsevier.com/retrieve/pii/S0959437X04001571

Burt, A., & Trivers, R. (2006). Genes in Conflict: The Biology of Selfish Genetic Elements. Harvard University Press.

Carballar-Lejarazú, R., Ogaugwu, C., Tushar, T., Kelsey, A., Pham, T. B., Murphy, J., Schmidt, H., Lee, Y., Lanzaro, G. C., & James, A. A. (2020). Next-generation gene drive for population modification of the malaria vector mosquito, Anopheles gambiae. Proceedings of the National Academy of Sciences, 117(37), 22805–22814. 10.1073/pnas.2010214117

Champer, J., Liu, J., Oh, S. Y., Reeves, R., Luthra, A., Oakes, N., Clark, A. G., & Messer, P. W. (2017). Reducing resistance allele formation in CRISPR gene drives (S. 150276). bioRxiv. 10.1101/150276

Chevalier, B. S., & Stoddard, B. L. (2001). Homing endonucleases: Structural and functional insight into the catalysts of intron/intein mobility. Nucleic Acids Research, 29(18), 3757–3774. 10.1093/nar/29.18.3757

Cotton, A. J., Földvári, M., Cotton, S., & Pomiankowski, A. (2014). Male eyespan size is associated with meiotic drive in wild stalk-eyed flies (Teleopsis dalmanni). Heredity, 112(4), 363–369. 10.1038/hdy.2013.131

Deredec, A., Burt, A., & Godfray, H. C. J. (2008). The Population Genetics of Using Homing Endonuclease Genes in Vector and Pest Management. Genetics, 179(4), 2013–2026. 10.1534/genetics.108.089037

Dong, Y., Simões, M. L., Marois, E., & Dimopoulos, G. (2018). CRISPR/Cas9 -mediated gene knockout of Anopheles gambiae FREP1 suppresses malaria parasite infection. PLOS Pathogens, 14(3), e1006898. 10.1371/journal.ppat.1006898

Drury, D. W., Dapper, A. L., Siniard, D. J., Zentner, G. E., & Wade, M. J. (2017). CRISPR/Cas9 gene drives in genetically variable and nonrandomly mating wild populations. Science Advances, 3(5), e1601910. 10.1126/sciadv.1601910

Esvelt, K. M., Smidler, A. L., Catteruccia, F., & Church, G. M. (2014). Concerning RNA-guided gene drives for the alteration of wild populations. eLife, 3, e03401. 10.7554/eLife.03401

Gantz, V. M., & Bier, E. (2015). The mutagenic chain reaction: A method for converting heterozygous to homozygous mutations. Science, 348(6233), 442–444. 10.1126/science.aaa5945

Gantz, V. M., Jasinskiene, N., Tatarenkova, O., Fazekas, A., Macias, V. M., Bier, E., & James, A. A. (2015). Highly efficient Cas9-mediated gene drive for population modification of the malaria vector mosquito Anopheles stephensi. Proceedings of the National Academy of Sciences, 112(49), E6736–E6743. 10.1073/pnas.1521077112

Grunwald, H. A., Gantz, V. M., Poplawski, G., Xu, X.-R. S., Bier, E., & Cooper, K. L. (2019). Super-Mendelian inheritance mediated by CRISPR–Cas9 in the female mouse germline. Nature, 566(7742), Article 7742. 10.1038/s41586-019-0875-2

Hammond, A., Galizi, R., Kyrou, K., Simoni, A., Siniscalchi, C., Katsanos, D., Gribble, M., Baker, D., Marois, E., Russell, S., Burt, A., Windbichler, N., Crisanti, A., & Nolan, T. (2016). A CRISPR-Cas9 gene drive system targeting female reproduction in the malaria mosquito vector Anopheles gambiae. Nature Biotechnology, 34(1), Article 1. 10.1038/nbt.3439

Hoermann, A., Habtewold, T., Selvaraj, P., Del Corsano, G., Capriotti, P., Inghilterra, M. G., Kebede, T. M., Christophides, G. K., & Windbichler, N. (2022). Gene drive mosquitoes can aid malaria elimination by retarding Plasmodium sporogonic development. Science Advances, 8(38), eabo1733. 10.1126/sciadv.abo1733

Hsu, P. D., Lander, E. S., & Zhang, F. (2014). Development and Applications of CRISPR-Cas9 for Genome Engineering. Cell, 157(6), 1262–1278. 10.1016/j.cell.2014.05.010

Jaenike, J. (1996). Sex-Ratio Meiotic Drive in the Drosophila quinaria Group. The American Naturalist, 148(2), 237–254. 10.1086/285923

Kyrou, K., Hammond, A. M., Galizi, R., Kranjc, N., Burt, A., Beaghton, A. K., Nolan, T., & Crisanti, A. (2018). A CRISPR–Cas9 gene drive targeting doublesex causes complete population suppression in caged Anopheles gambiae mosquitoes. Nature Biotechnology, 36(11), Article 11. 10.1038/nbt.4245

Lambrechts, L., Koella, J. C., & Boëte, C. (2008). Can transgenic mosquitoes afford the fitness cost? Trends in Parasitology, 24(1), 4–7. 10.1016/j.pt.2007.09.009

Lande, R., & Wilkinson, G. S. (1999). Models of sex-ratio meiotic drive and sexual selection in stalk-eyed flies. Genetics Research, 74(3), 245–253. 10.1017/S0016672399004218

Lehmann, L., Keller, L. F., & Kokko, H. (2007). Mate choice evolution, dominance effects, and the maintenance of genetic variation. Journal of Theoretical Biology, 244(2), 282–295. 10.1016/j.jtbi.2006.07.033

Lehmann, T., Hawley, W. A., Grebert, H., & Collins, F. H. (1998). The effective population size of Anopheles gambiae in Kenya: Implications for population structure. Molecular Biology and Evolution, 15(3), 264–276. 10.1093/oxfordjournals.molbev.a025923

Lenington, S. (1991). The t Complex: A Story of Genes, Behavior, and Populations. In P. J. B. Slater, J. S. Rosenblatt, C. Beer, & M. Milinski (Hrsg.), Advances in the Study of Behavior (Bd. 20, S. 51–86). Academic Press.

Li, M., Yang, T., Kandul, N. P., Bui, M., Gamez, S., Raban, R., Bennett, J., Sánchez C, H. M., Lanzaro, G. C., Schmidt, H., Lee, Y., Marshall, J. M., & Akbari, O. S. (2020). Development of a confinable gene drive system in the human disease vector Aedes aegypti. eLife, 9, e51701. 10.7554/eLife.51701

Lindholm, A. K., Dyer, K. A., Firman, R. C., Fishman, L., Forstmeier, W., Holman, L., Johannesson, H., Knief, U., Kokko, H., Larracuente, A. M., Manser, A., Montchamp- Moreau, C., Petrosyan, V. G., Pomiankowski, A., Presgraves, D. C., Safronova, L. D., Sutter, A., Unckless, R. L., Verspoor, R. L., … Price, T. A. R. (2016). The Ecology and Evolutionary Dynamics of Meiotic Drive. Trends in Ecology & Evolution, 31(4), 315–326. 10.1016/j.tree.2016.02.001

Lyttle, T. W. (1993). Cheaters sometimes prosper: Distortion of mendelian segregation by meiotic drive. Trends in Genetics, 9(6), 205–210. 10.1016/0168-9525(93)90120-7

Manser, A., König, B., & Lindholm, A. K. (2020). Polyandry blocks gene drive in a wild house mouse population. Nature Communications, 11(1), Article 1. 10.1038/s41467-020-18967-8

Manser, A., Lindholm, A. K., & Weissing, F. J. (2017). The evolution of costly mate choice against segregation distorters. Evolution, 71(12), 2817–2828. 10.1111/evo.13376

Meccariello, A., Hou, S., Davydova, S., Fawcett, J. D., Siddall, A., Leftwich, P. T., Krsticevic, F., Papathanos, P. A., & Windbichler, N. (2024). Gene drive and genetic sex conversion in the global agricultural pest Ceratitis capitata. Nature Communications, 15(1), 372. 10.1038/s41467-023-44399-1

Mendel, G. (1865). Verhandlungen des naturforschenden Vereins. In *Versuche über Pflanzenhybriden* (Bd. 4, S. 3–47). Im Verlage des Vereins.

Moro, D., Byrne, M., Kennedy, M., Campbell, S., & Tizard, M. (2018). Identifying knowledge gaps for gene drive research to control invasive animal species: The next CRISPR step. Global Ecology and Conservation, 13, e00363. 10.1016/j.gecco.2017.e00363

Nabutanyi, P., & Wittmann, M. J. (2021). Models for Eco-Evolutionary Extinction Vortices under Balancing Selection. The American Naturalist, 197(3), 336–350. 10.1086/712805

Pomiankowski, A. (1987). The costs of choice in sexual selection. Journal of Theoretical Biology, 128(2), 195–218. 10.1016/S0022-5193(87)80169-8

Price, T. A. R., & Wedell, N. (2008). Selfish genetic elements and sexual selection: Their impact on male fertility. Genetica, 134(1), 99–111. 10.1007/s10709-008-9253-y

Price, T. A. R., Windbichler, N., Unckless, R. L., Sutter, A., Runge, J.-N., Ross, P. A., Pomiankowski, A., Nuckolls, N. L., Montchamp-Moreau, C., Mideo, N., Martin, O. Y., Manser, A., Legros, M., Larracuente, A. M., Holman, L., Godwin, J., Gemmell, N., Courret, C., Buchman, A., … Lindholm, A. K. (2020). Resistance to natural and synthetic gene drive systems. Journal of Evolutionary Biology, 33(10), 1345–1360. 10.1111/jeb.13693

R Core Team. (2022). *R: A Language and Environment for Statistical Computing* [Software]. R Foundation for Statistical Computing. https://www.R-project.org/

Ran, F. A., Hsu, P. D., Wright, J., Agarwala, V., Scott, D. A., & Zhang, F. (2013). Genome engineering using the CRISPR-Cas9 system. Nature Protocols, 8(11), 2281–2308. 10.1038/nprot.2013.143

Rode, N. O., Estoup, A., Bourguet, D., Courtier-Orgogozo, V., & Débarre, F. (2019). Population management using gene drive: Molecular design, models of spread dynamics and assessment of ecological risks. Conservation Genetics, 20(4), 671–690. 10.1007/s10592-019-01165-5

Sandler, L., & Novitski, E. (1957). Meiotic Drive as an Evolutionary Force. The American Naturalist, 91(857), 105–110. 10.1086/281969

Silver, L. M. (1993). The peculiar journey of a selfish chromosome: Mouse t haplotypes and meiotic drive. Trends in Genetics, 9(7), 250–254. 10.1016/0168-9525(93)90090-5

Simoni, A., Hammond, A. M., Beaghton, A. K., Galizi, R., Taxiarchi, C., Kyrou, K., Meacci, D., Gribble, M., Morselli, G., Burt, A., Nolan, T., & Crisanti, A. (2020). A male-biased sex-distorter gene drive for the human malaria vector Anopheles gambiae. Nature Biotechnology, 38(9), 1054–1060. 10.1038/s41587-020-0508-1

Stoddard, B. L. (2005). Homing endonuclease structure and function. Quarterly Reviews of Biophysics, 38(1), 49–95. 10.1017/S0033583505004063

Sutter, A., & Lindholm, A. K. (2016). No evidence for female discrimination against male house mice carrying a selfish genetic element. Current Zoology, 62(6), 675–685. 10.1093/cz/zow063

Unckless, R. L., Clark, A. G., & Messer, P. W. (2017). Evolution of Resistance Against CRISPR/Cas9 Gene Drive. Genetics, 205(2), 827–841. 10.1534/genetics.116.197285

Unckless, R. L., Messer, P. W., Connallon, T., & Clark, A. G. (2015). Modeling the Manipulation of Natural Populations by the Mutagenic Chain Reaction. Genetics, 201(2), 425–431. 10.1534/genetics.115.177592

Verma, P., Reeves, R. G., Simon, S., Otto, M., & Gokhale, C. S. (2023). The Effect of Mating Complexity on Gene Drive Dynamics. The American Naturalist, 201(1), E1–E22. 10.1086/722157

Wang, S., & Jacobs-Lorena, M. (2013). Genetic approaches to interfere with malaria transmission by vector mosquitoes. Trends in Biotechnology, 31(3), 185–193. 10.1016/j.tibtech.2013.01.001

Wedell, N. (2013). The dynamic relationship between polyandry and selfish genetic elements. Philosophical Transactions of the Royal Society B: Biological Sciences, 368(1613), 20120049. 10.1098/rstb.2012.0049

Wedell, N., Price, T. a. R., & Lindholm, A. K. (2019). Gene drive: Progress and prospects. Proceedings of the Royal Society B: Biological Sciences, 286(1917), 20192709. 10.1098/rspb.2019.2709

WHO. (2020, März 2). Vector-borne diseases. https://www.who.int/news-room/fact-sheets/detail/vector-borne-diseases

WHO. (2023). *World malaria report* 2023. https://www.who.int/teams/global-malaria-programme/reports/world-malaria-report-2023

Wilkinson, G. S., Presgraves, D. C., & Crymes, L. (1998). Male eye span in stalk-eyed flies indicates genetic quality by meiotic drive suppression. Nature, 391(6664), 276–279. 10.1038/34640

Yadav, A. K., Butler, C., Yamamoto, A., Patil, A. A., Lloyd, A. L., & Scott, M. J. (2023). CRISPR/Cas9-based split homing gene drive targeting doublesex for population suppression of the global fruit pest Drosophila suzukii. Proceedings of the National Academy of Sciences, 120(25), e2301525120. 10.1073/pnas.2301525120

