## Supplementary Materials for "Mate choice against gene drives leads to evolutionary rescue"

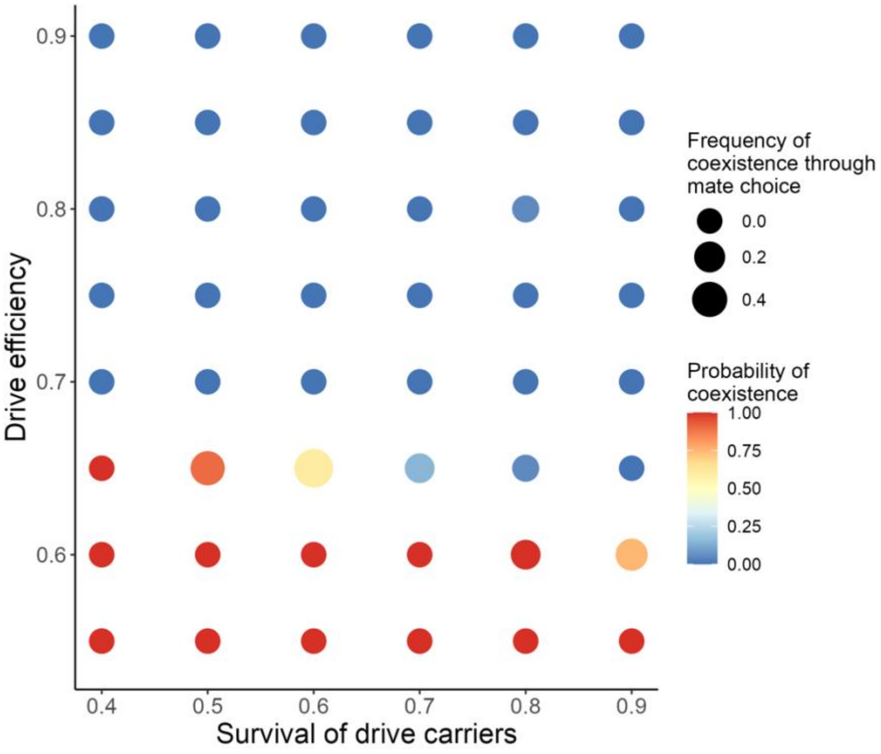


**Figure S1. Effects of drive efficiency, survival of drive carriers and mate choice on the probability of allele coexistence for recessive drives.** This figure illustrates the probability of coexistence and the frequency of coexistence due to mate choice under conditions where the gene drive was fully recessive ($h$ = 0), across various levels of drive efficiency and survival of drive carriers (20 replicates per scenario). Coexistence is defined as the scenario where both wild-type and gene-drive alleles maintain frequencies between 0.01 and 0.99 at the 200 generation. The probability of coexistence is calculated as the average occurrence of coexistence across the 20 replicates (represented by circle colours). The frequency of coexistence due to mate choice is determined by comparing the probability of coexistence in simulations with evolving mating preferences to those with random mating (represented by circle size, with larger circles indicating a higher frequency of coexistence attributed to the evolution of female preferences). All other parameter values are as specified in Table 1.

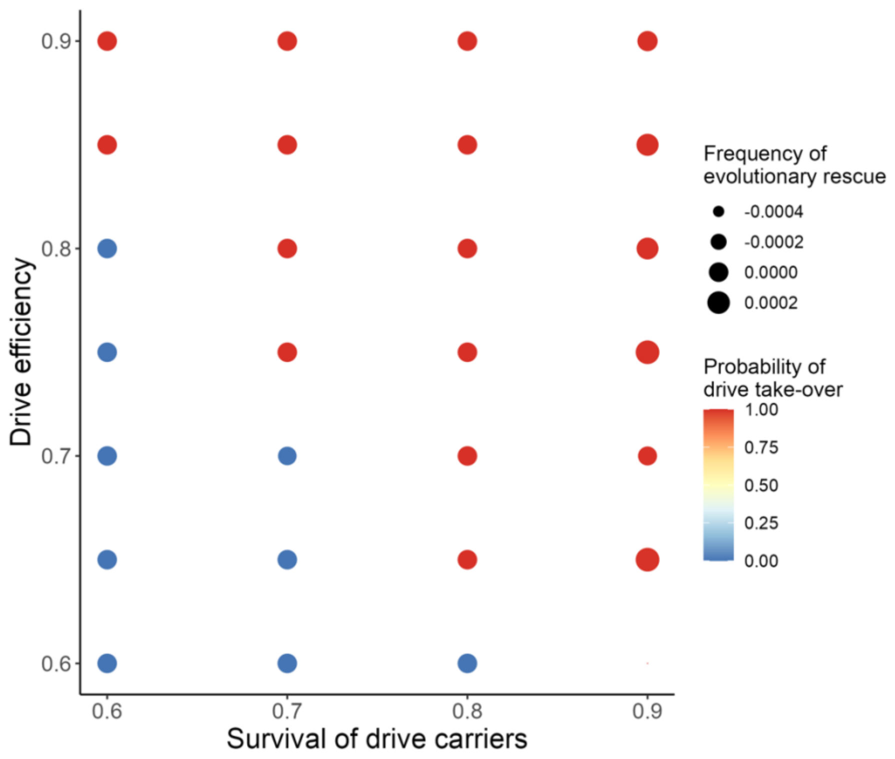


**Figure S2. Effect of small initial variance in mating preferences on the probability of gene drive take-over and the occurrence of evolutionary rescue.** The probability of drive take-over and the frequency of evolutionary rescue are shown under conditions where the initial variance in mating preferences was low ($\sigma_0 = 0.1$), across various levels of drive efficiency and survival of drive carriers (20 replicates per scenario). The probability of drive take-over is measured as the average frequency of the drive allele at generation 200 in scenarios with evolving mating preferences (represented by circle colour). The frequency of evolutionary rescue is calculated as the difference in the probability of drive take-over between simulations that include the evolution of mating preferences and those with random mating (represented by circle size, with larger circles indicating a higher occurrence of evolutionary rescue due to mate choice). Here, the gene drive was fully dominant ($h$ = 1). All other parameter values are as specified in Table 1.

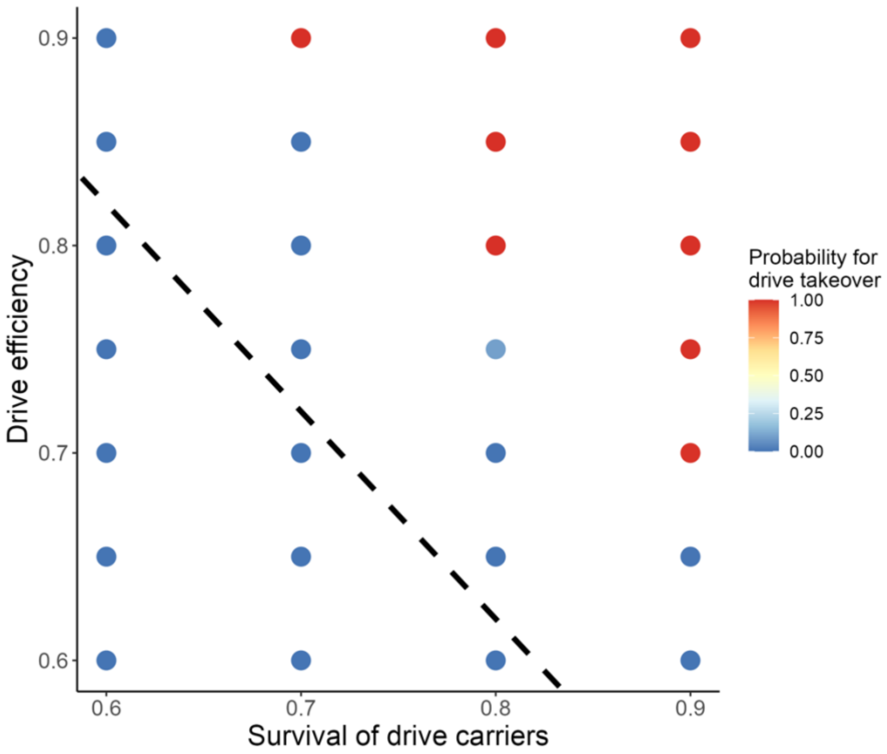


**Figure S3. Impact of initial mating preferences for wild-type males on the probability of gene drive take-over.** This figure shows the probability of drive take-over when females initially preferred wild-type males on average ($\mu_0$ = 1), across various levels of drive efficiency and survival of drive carriers (20 replicates per scenario). The probability of drive take-over is measured as the average frequency of the drive allele at generation 200 in scenarios with evolving mating preferences (represented by circle colours). The reference line indicates the threshold between drive take-over and loss of the drive in simulations without any mating preferences (as shown in Fig. 5). Here, the gene drive was fully dominant ($h$ = 1). All other parameter values are as specified in Table 1.

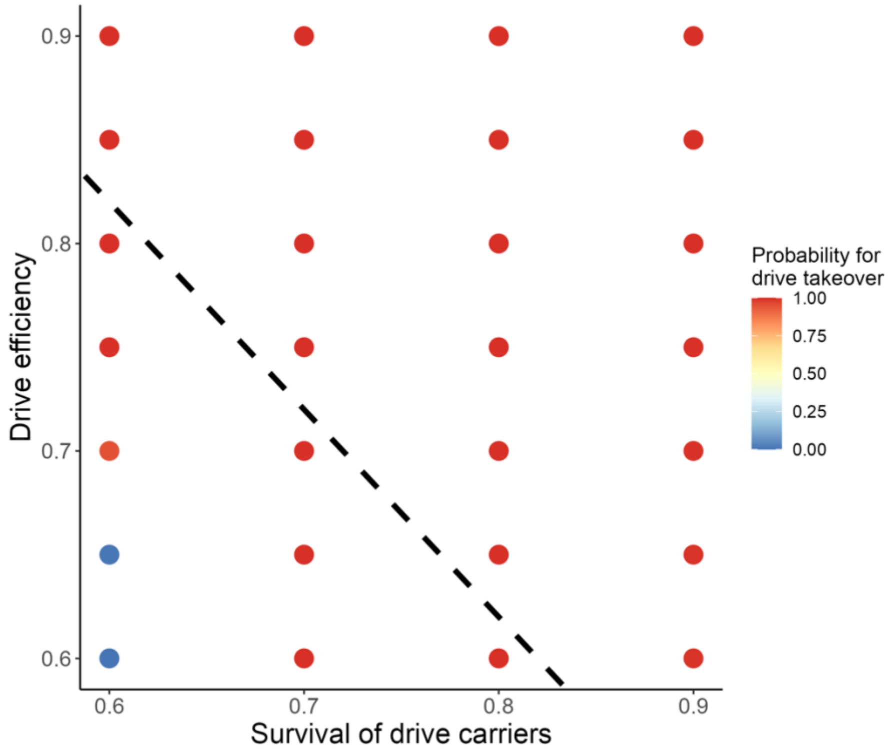


**Figure S4. Impact of initial mating preferences for drive carriers on the probability of gene drive take-over.** This figure shows the probability of drive take-over when females initially preferred drive carriers on average ($\mu_0 = -1$), across various levels of drive efficiency and survival of drive carriers (20 replicates per scenario). The probability of drive take-over is measured as the average frequency of the drive allele at generation 200 in scenarios with evolving mating preferences (represented by circle colours). The reference line indicates the threshold between drive take-over and loss of the drive in simulations without any mating preferences (as shown in Fig. S5). Here, the gene drive was fully dominant ($h$ = 1). All other parameter values are as specified in Table 1.

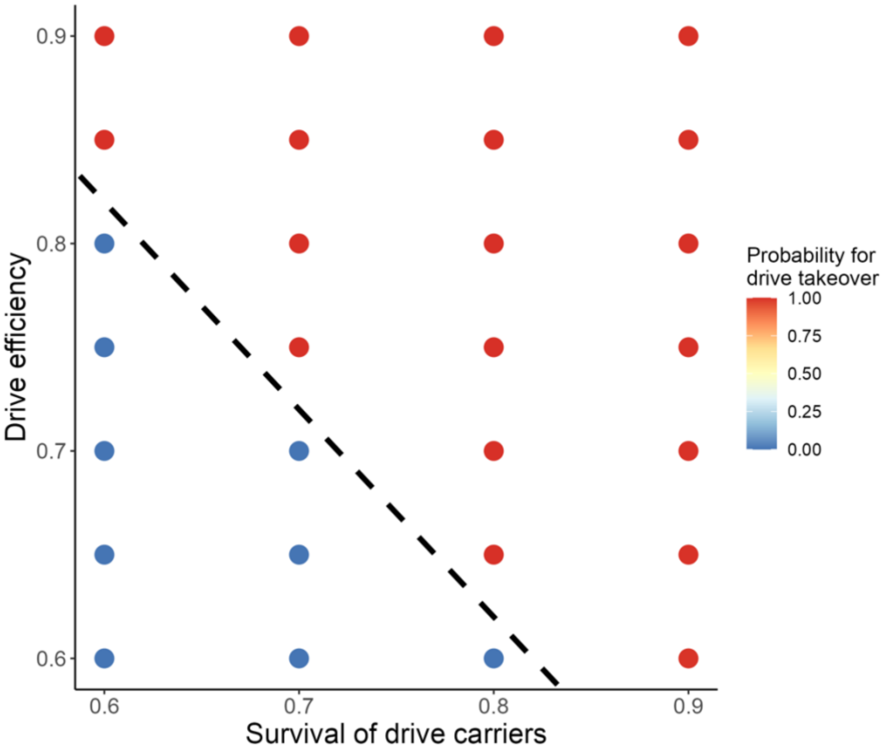


**Figure S5. Probability of gene drive take-over under random mating conditions.** This figure shows the probability of drive take-over across various levels of drive efficiency and survival of drive carriers with random mating (i.e., no evolving mating preferences). The probability of drive take-over is measured as the average frequency of the drive allele at generation 200 under random mating conditions across 20 replicates (represented by circle colours). The reference line indicates the threshold between drive take-over (red) and loss of the drive (blue) within these simulations. This reference line is used in Figs. S3 and S4 to facilitate comparison between evolutionary outcomes of simulations with initial preferences for wild-type males and simulations with initial preferences for drive carriers. Here, the gene drive was fully dominant ($h$ = 1)**.** All parameter values are as specified in Table 1._
